# Altered liver metabolism post-wean abolishes efficacy of vitamin D for breast cancer prevention in a mouse model

**DOI:** 10.1101/2024.05.28.596304

**Authors:** Sarah M Bernhardt, Michelle K Ozaki, Courtney Betts, Lisa A Bleyle, Andrea E DeBarber, Jaime Fornetti, Abigail L Liberty, Elise De Wilde, Yi Zhang, Zheng Xia, Pepper Schedin

## Abstract

Young women have increased risk of vitamin D deficiency, which may increase breast cancer incidence. Here, we assessed the anti-cancer efficacy of vitamin D in mouse models of young-onset breast cancer. In never-pregnant mice, vitamin D supplementation increased serum 25(OH)D and hepatic 1,25(OH)_2_D_3_, reduced tumor size, and associated with anti-tumor immunity. These anti-tumor effects were not replicated in a mouse model of postpartum breast cancer, where hepatic metabolism of vitamin D was suppressed post-wean, which resulted in deficient serum 25(OH)D and reduced hepatic 1,25(OH)_2_D_3_. Treatment with active 1,25(OH)_2_D_3_ induced hypercalcemia exclusively in post-wean mice, highlighting metabolic imbalance post-wean. RNAseq revealed suppressed CYP450 expression postpartum. In sum, we provide evidence that vitamin D anti-tumor activity is mediated through immunomodulatory mechanisms and is ineffective in the post-wean window due to altered hepatic metabolism. These findings have implications for suppressed xenobiotic metabolism in postpartum women beyond vitamin D.

**Statement of Significance:** In a rodent model of postpartum breast cancer, weaning suppresses hepatic CYP450 activity and renders vitamin D supplementation ineffective, with implications for xenobiotic drug efficacy and safety. A tailored approach to therapy based on reproductive history is crucial for young breast cancer patients, and for healthcare strategies for postpartum women.

## Introduction

Women have an increased risk of developing breast cancer within 10 years of a completed pregnancy (1, 2). Breast cancers that arise in this postpartum period are referred to as postpartum breast cancer (PPBC) (3, 4). PPBC is associated with increased rates of metastasis and poorer survival outcomes (3, 5–9). PPBC accounts for ∼19% of all new breast cancer globally, and ∼30-50% of all young-onset breast cancer (10). This makes PPBC a major health problem for young women worldwide. In rodent models, the physiologic event of postpartum mammary gland involution is a dominant mediator of PPBC progression, and thus a potential target for prevention strategies (11–13). Mammary gland involution occurs after parturition in the absence of lactation, or with weaning. During involution, the mammary epithelial cells that were expanded for milk production are eliminated through a well-characterized, developmentally regulated process (14, 15). In both rodents and humans, the involution mammary microenvironment shares similarities with wound-healing, tumor desmoplasia, and inflammation (16–21). Further, postpartum involution is causal to PPBC progression in rodent models (11–13, 19, 22). Postpartum involution also independently associates with increased rates of breast cancer metastasis and poor prognostic tumor gene signatures observed in young women (3, 5–7, 23).

The involuting mammary gland is characterized by an immunosuppressive microenvironment (18, 24). While overall immune cell abundance increases in the breast during involution, there is an influx of immature and M2-skewed myeloid cells (17, 25). In rodents, there is functional evidence for active T-cell suppression (24). Consequently, targeting mammary gland involution with immunotherapeutic agents that act on immature myeloid cells is a potential prevention strategy for PPBC (26, 27). As proof-of-principal, administration of non-steroidal anti-inflammatory agents (NSAIDs) during involution significantly reduced PPBC incidence and progression in mice (12, 13, 19). Specifically, NSAID treatment in murine models of PPBC associated with reduced abundance of immature myeloid cells and a shift from a pro-tumor Th2 to an antitumor Th1 immune milieu (12). However, NSAIDs administered at the time of weaning did not completely mitigate the pro-tumor attributes of mammary gland involution in murine models of PPBC (12, 13). This suggests that additional targeting of the inflammatory milieu that is present post-wean might improve efficacy.

Vitamin D is an immunomodulatory agent that has an excellent safety profile (28). In women, increased serum vitamin D, defined as 25(OH)D, associates with decreased risk of breast cancer (29–31). In women with breast cancer, increased serum 25(OH)D associates with reduced risk of mortality (32). One mechanism through which vitamin D might reduce breast cancer risk is by mitigating the immunosuppressive activity of immature myeloid cells. The immune-modulatory effects of vitamin D are mediated by its active form, 1,25-dihydroxyvitamin D_3_ [1,25(OH)_2_D_3_], which binds the vitamin D receptor (VDR) (33). Importantly, immature myeloid cells express VDR, and are identified as direct targets of 1,25(OH)_2_D_3_ (34). Of note, the postpartum window is a period of vitamin D deficiency, due to the elevated demand for vitamin D and calcium during pregnancy and lactation. Further, the postpartum period is characterized by immunosuppression in the mammary gland (18, 24). In the US, vitamin D deficiency is observed in up to 72% of postpartum women (35, 36). This deficiency might further exacerbate breast cancer risk in an already vulnerable postpartum period. Together, these previous studies suggests that vitamin D supplementation at time of weaning might have efficacy against PPBC through promotion of anti-tumor activity (37).

Currently, the effect of vitamin D treatment on myeloid cell populations in breast cancer remains under-studied. Whether vitamin D supplementation is more effective when targeted to an immune-suppressive, vitamin D-deficient window, such as postpartum mammary gland involution, has not been addressed. Here, we investigated the anti-cancer potential of vitamin D in mouse models of young-onset breast cancer. Our objective was to explore the underlying mechanisms that drive vitamin D’s anti-cancer effects. We found vitamin D supplementation reduced mammary tumor growth in never-pregnant mice, likely through promotion of macrophage maturation and relief of T-cell suppression. However, vitamin D supplementation of involution mice failed to reduce PPBC growth. We determined that suppressed vitamin D metabolism in the post-wean liver abolished the anti-tumor efficacy of vitamin D supplementation. This abolition of efficacy was mediated through altered cytochrome P450 gene expression. We attempted to circumvent this impaired metabolism through administering the active form of vitamin D. However, this proved fatal in postpartum mice due to the altered calcium regulation associated with weaning. These findings support vitamin D as an immunomodulatory, anti-cancer agent for breast cancers that do not arise postpartum. However, our data identify the post-wean period as a unique metabolic window wherein vitamin D supplementation is ineffective. These findings broaden our understanding of how reproductive stage regulates liver cytochrome P450 enzyme expression, with implications for the metabolism of endocrine and chemotherapeutic agents in PPBC patients. Together, our results highlight the necessity for additional research to address whether suppressed P450 metabolic activity postpartum puts women at risk for reduced xenobiotic drug activity as well as increased drug toxicity.

## Results

### Vitamin D reduces tumor burden in nulliparous mice, and associates with an anti-tumor immune response

We first established a murine model of dietary vitamin D deficiency and supplementation reflecting levels observed in humans. To achieve this, adult never-pregnant (nulliparous; nul) female BALB/c mice were fed diets deficient (Low; 25IU/kg) or supplemented (High; 4,000IU/kg) with vitamin D for 5 weeks. Blood was collected at study end and serum vitamin D, defined by circulating concentrations of 25(OH)D, were assessed. Nulliparous mice fed a vitamin D-supplemented diet showed a >2-fold increase in serum 25(OH)D (67.4±8.1nmol/L) compared to mice fed a vitamin D-deficient diet (28.7±11.7nmol/L, p<0.01; Figure 1A). These serum levels model human vitamin D deficiency and supplementation, respectively, as defined by the Institute of Medicine guidelines (IOM, USA). The IOM vitamin D deficiency and supplementation levels are depicted by the dashed lines in Figure 1A (38).

**Figure 1:**
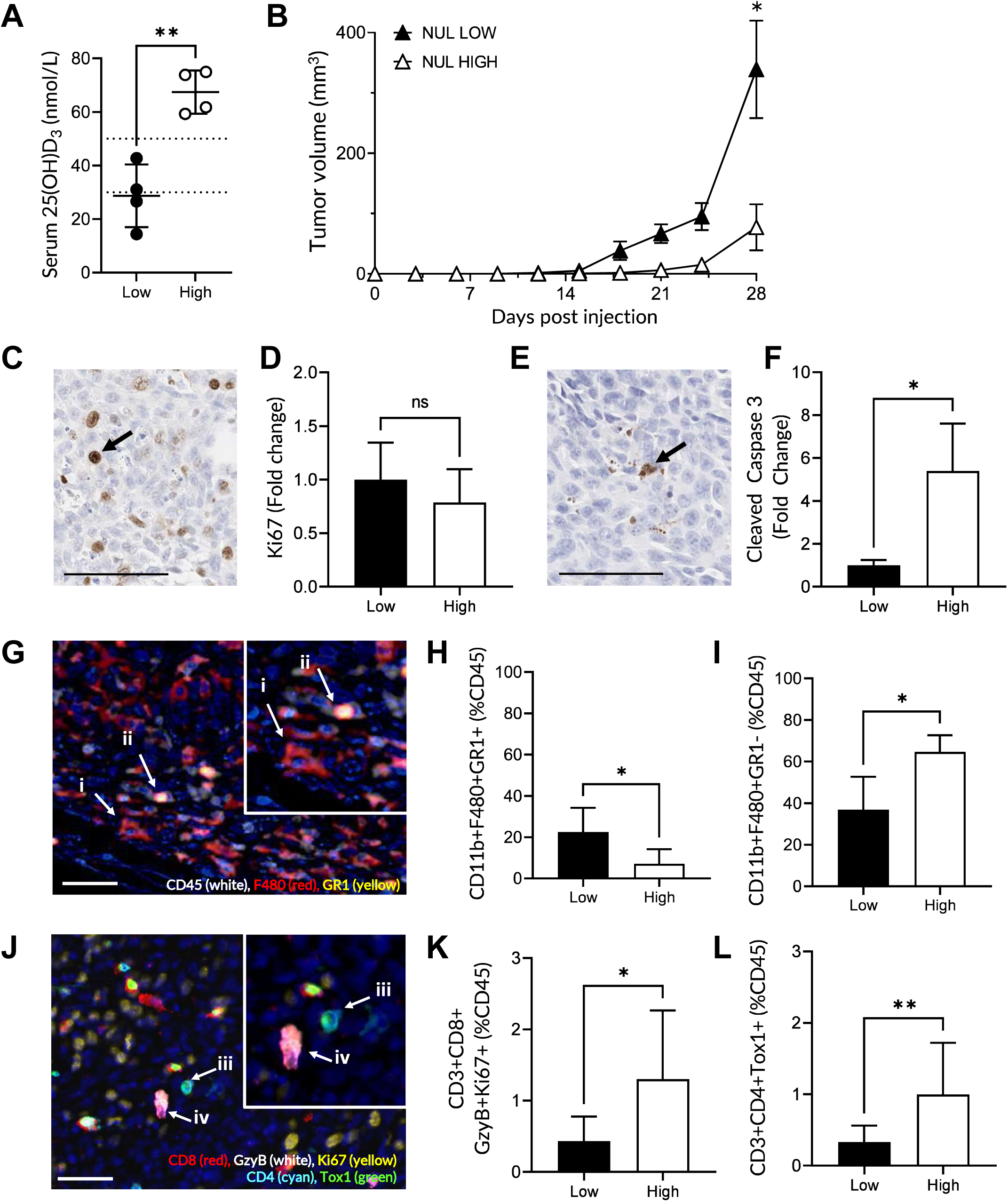
Vitamin D supplementation inhibits tumor growth in nulliparous mice, likely through promotion of macrophage maturation and relief of T-cell suppression. (A) Serum 25(OH)D in nulliparous mice fed diets deficient (Low, 25IU/kg) or supplemented (High, 4,000IU/kg) for vitamin D, quantified by LC-MS/MS. Dashed lines represent vitamin D deficiency (<30nmol/L) and sufficiency (>50nmol/L). (B-L) Mouse mammary tumor model in nulliparous (Nul) mice, using vitamin D deficient (Nul Low, n=9 mice), or supplemented (Nul High, n=5) diets. (B) Tumor growth over 4 weeks. (C) Representative staining of Ki67. (D) Quantification of tumor Ki67 staining. (E) Representative staining of cleaved caspase 3 (CC3). (F) Quantification of tumor CC3 staining in tumors. (G) Representative multiplex immunohistochemistry (IHC) staining of CD45^+^ (white), F480^+^ (red) and GR1^+^ (yellow). Quantification of abundance of (H) immature myeloid cells (CD45^+^CD11b^+^F480^+^GR1^+^) identified by arrows (ii); and (I) mature macrophages (CD45^+^CD11b^+^F480^+^GR1^-^) represented by arrows (i). (J) Representative multiplex IHC staining of CD8^+^ (red), GzyB^+^ (white), Ki67^+^ (yellow), CD4^+^ (cyan), Tox1^+^ (green). (K) Quantification of abundance of activated CD8^+^ T-cells (CD45^+^CD3^+^CD8^+^GzyB^+^Ki67^+^) identified by arrows (iv); and (L) activated CD4^+^ T-cells (CD45^+^CD3^+^CD4^+^Tox1^+^) identified by arrows (iii). Data are mean±Stdev. P-values are presented, determined through two-tailed T-test. ns= not significant; * p<0.05; **p<0.01.

We next assessed if vitamin D deficiency and supplementation affects mammary tumor growth in nulliparous hosts. Mouse mammary tumor cells (D2A1, 20,000 cells in PBS, 20µL) were injected into the fourth (inguinal) mammary fat-pads. Tumor growth was monitored for 4 weeks. In supplemented mice, elevated serum vitamin D associated with a 3-fold reduction in tumor growth (p=0.03), compared to vitamin D deficient mice (Figure 1B). Given the known anti-proliferative and pro-apoptotic roles of vitamin D (37), we next assessed tumor proliferation (Ki67) and apoptosis (cleaved caspase 3; CC3) through immunohistochemistry (IHC). We found that vitamin D supplementation did not affect tumor proliferation (p=0.33; Figure 1C,D). In contrast, vitamin D supplementation increased tumor apoptosis 5.3-fold (p=0.04; Figure 1E,F).

Increased tumor cell death with vitamin D supplementation could implicate increased T cell-mediated cytotoxicity. For example, mature myeloid cells are reported to promote T-cell activation and tumor cell killing via apoptotic pathways marked by CC3. In contrast, their immature precursors are implicated in T-cell suppression. Based on the observation that vitamin D supplementation increased CC3-mediated cell death, we assessed tumor-associated myeloid and T-cell abundance and phenotype using multiplex IHC. We found the total number of immature myeloid cells (defined as CD45^+^CD11b^+^F480^+^GR1^+^) was significantly reduced in tumors collected from vitamin D-supplemented mice (7.2±7% of total CD45^+^ immune cells), compared to vitamin D-deficient mice (22.5±11.7%; p=0.04; Figure 1G,H). In contrast, in vitamin D-supplemented group tumors, the number of mature macrophages (CD45^+^CD11b^+^F480^+^GR1^-^) were significantly elevated (64.4±8.0%) compared to tumors from vitamin D-deficient mice (36.9±15.8%; p=0.01; Figure 1G,I). These results suggest a role for vitamin D in the maturation of tumor-associated immature myeloid cells.

Immature myeloid cells play a role in T-cell suppression (39). Given this, we anticipated increased abundance of activated T-cells with vitamin D supplementation. To assess T-cell activation, we used Granzyme B (GzyB), Ki67, and TOX1 as markers of T-cell activation/exhaustion. GzyB is secreted by cytotoxic T-cells for tumor cell elimination via caspase-dependent and independent pathways (40). Ki67 is a marker of T-cell proliferation. TOX1 is a marker of T-cell exhaustion on cytotoxic CD8 T cells, whereas increased TOX1 expression on CD4 cells is associated with activated T-helper cells. We found that vitamin D supplementation increased abundance of intratumoral activated cytotoxic CD8^+^ T-cells (CD45^+^CD3^+^CD8^+^GzyB^+^Ki67^+^) ∼3-fold (1.3±1.0% of total CD45^+^ vs deficient, 0.4±0.3; p=0.013; Figure 1J,K), and increased activated helper CD4^+^ T-cells ∼3-fold (CD45^+^CD3^+^CD4^+^TOX1^+^) compared to tumors collected from vitamin D-deficient mice (supplemented, 1.0±0.7% vs deficient, 0.3±0.2; p<0.01; Figure 1J,L).

### Weaning-induced mammary gland involution is a rational target for vitamin D-based intervention

The post-wean period is a transient window characterized by vitamin D deficiency (35, 36) and local mammary gland immune suppression (18, 24). Our findings suggests that the postpartum window might be highly responsive to vitamin D supplementation (37). In our postpartum mouse model, we first confirmed the extent of vitamin D deficiency in the post-wean window. In mice fed adequate levels of vitamin D, we found baseline serum 25(OH)D was reduced in involution mice (27.6±4.6nmol/L), at a concentration defined as ‘vitamin D deficient’ (38), compared to nulliparous mice (38.9±10.3nmol/L; p=0.03; Figure 2A). As vitamin D functions to regulate calcium status, with one mechanism through mobilization of calcium from the bone (41), we also assessed serum calcium and bone mineral density in mice following weaning. Serum calcium was significantly elevated in involution mice (9.58±2.04mg/dL) compared to nulliparous mice (7.39±1.27mg/dL; p=0.04; Figure 2B). In parallel, bone mineral density was significantly reduced in involution mice (73.3±6.8%), compared to nulliparous mice (91.7±12.0%, p<0.001; Figure 2C). Thus, this mouse model appears to reflect the vitamin D, calcium, and bone mineral density biology observed in recently pregnant and lactating women.

**Figure 2:**
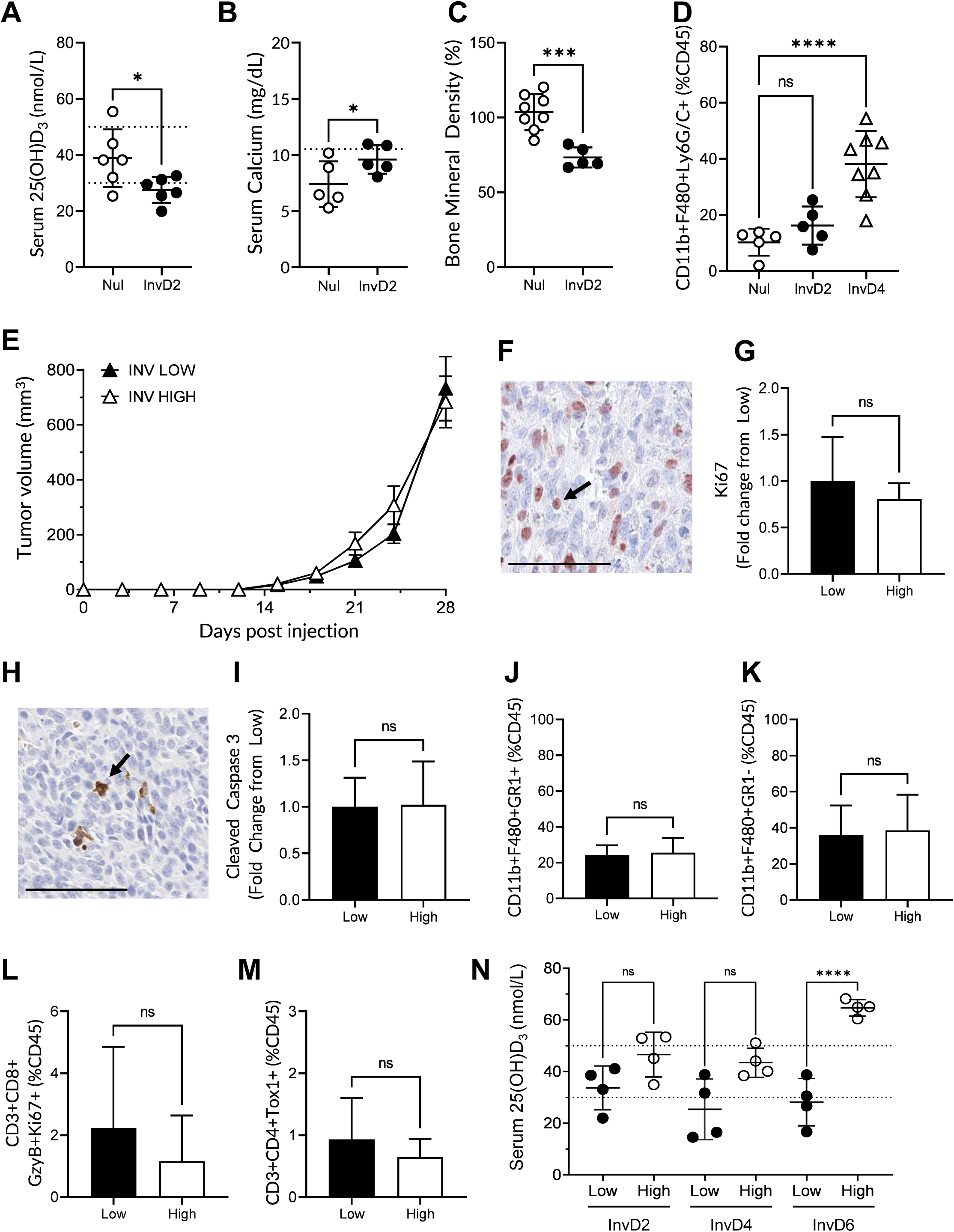
Vitamin D supplementation is ineffective at reducing tumor growth in the postpartum period. (A) Serum 25(OH)D. Dotted lines represent human vitamin D deficiency (30nmol/L) and sufficiency (50nmol/L). (B) Serum calcium in nulliparous (Nul) and involution day 2 (InvD2) mice fed a vitamin D adequate diet (1,000IU/kg). (C) Bone mineral density in mice fed vitamin D supplemented diets (4,000IU/kg). (D) Flow cytometry of digested mammary glands collected from Nul, InvD2 and InvD4 mice quantifying immature myeloid cell (CD45^+^CD11b^+^F480^+^Ly6G/C+) abundance. (E-N) Mouse mammary postpartum tumor model under vitamin D deficient (Inv Low, n=12 mice), or supplemented (Inv High, n=6) diets. (E) Tumor growth over 4 weeks. (F) Representative staining of Ki67 (marker of proliferation) and (G) quantification of Ki67 in tumors at study end. (H) Representative staining of cleaved caspase 3 (CC3) (marker of apoptosis) and (I) quantification of CC3 staining in tumors. Quantification of multiplex immunohistochemistry (IHC) for abundance of (J) immature myeloid cells (CD45^+^CD11b^+^F480^+^GR1^+^); (K) mature macrophages (CD45^+^CD11b^+^F480^+^GR1^-^); (L) activated CD8^+^ T-cells (CD45^+^CD3^+^CD8^+^ GzyB^+^Ki67^+^); and (M) activated CD4^+^ T-cells (CD45^+^CD3^+^CD4^+^Tox1^+^). (N) Serum 25(OH)D in InvD2, InvD4, InvD6 mice fed diets deficient or supplemented for vitamin D, quantified through LC-MS/MS. Data are mean±Stdev. P-values are presented, determined through either T-test (2 group comparison) or ANOVA (multiple group comparison) with multiple comparisons. ns= not significant; * p<0.05; **p<0.01; ***p<0.001, ****p<0.0001.

We next assessed if the proposed targets of vitamin D efficacy, immature myeloid cells, were increased in the normal, involuting murine mammary gland. Immature myeloid cell abundance was significantly increased in involution day 4 mammary glands (38.1±11.8% of total CD45^+^ immune cells), compared to nulliparous (10.27±4.8%; p<0.001; Figure 2D). Thus, with evidence of vitamin D deficiency and increased abundance of the vitamin D immune target during involution, we rationalized that vitamin D supplementation during involution would be highly efficacious at reducing tumor growth in this model of PPBC.

### Vitamin D supplementation does not affect tumor growth or alter the immune milieu in the post-wean period

Vitamin D deficiency and supplementation were established during the post-wean window through dietary intake, as described above. D2A1 mouse mammary tumor cells were injected into the fourth inguinal fat-pads of mice at two days post-wean (involution day 2, InvD2). Surprisingly, vitamin D supplementation did not reduce tumor growth in involution mice (final volume 682.9±94.0mm^3^), compared to vitamin D-deficient mice (731.8±116.7mm^3^; p=0.79; Figure 2E). Consistent with a lack of difference in tumor size, we found no differences in tumor cell proliferation (p=0.41; Figure 2F,G) or apoptosis (p=0.97; Figure 2H,I) between vitamin D-deficient and supplemented mice. Similarly, multiplex IHC analysis revealed no differences in the abundance of mature macrophages (Figure 2J), immature myeloid cells (Figure 2K), nor in the activation of CD8^+^ (Figure 2L) or CD4^+^ (Figure 2M) T-cells.

To begin to understand the lack of efficacy, we returned to our non-tumor mouse model. We assessed if circulating concentrations of vitamin D metabolite, 25(OH)D differ by reproductive state. In contrast to that observed in nulliparous mice, who responded to vitamin D supplementation with appropriately elevated physiologic serum 25(OH)D (Figure 1A), restoration of serum 25(OH)D levels was delayed in postpartum mice. Serum collected on involution day 2 (supplemented 46.6±8.7nmol/L vs deficient 33.7±8.5nmol/L, p>0.05) and involution day 4 (supplemented 43.4±5.6nmol/L vs deficient 25.4±11.8nmol/L, p>0.05) did not exhibit differences in 25(OH)D between vitamin D-deficient and supplemented mice (Figure 2N). By involution day 6, vitamin D supplementation showed a >2-fold increase in serum 25(OH)D (64.7±3.2nmol/L) compared to deficient mice (28.2±9.1nmol/L, p<0.0001)(Figure 2N). Together, these findings highlight a transient, post-wean period where vitamin D supplementation is unable to restore serum 25(OH)D to sufficiency. Based on this observation, we next evaluated whether the primary enzymes involved in the metabolism of dietary vitamin D differed between nulliparous and involution mice.

### Vitamin D metabolism is impaired in the post-wean liver

Dietary vitamin D is biologically inactive and requires two hydroxylation steps to produce its active form (Figure 3A)(42). Dietary vitamin D (Vitamin D_3_) is hydroxylated in the liver by 25-hydroxylase (Cyp2r1) to produce 25(OH)D, the circulating form of vitamin D, and further metabolized by 1α-hydroxylase (Cyp27b1) to yield the active form (1,25(OH)_2_D_3_). We next identified potential factors that might contribute to reduced serum 25(OH)D in the postpartum window. We assessed gene expression of these enzymes in livers collected from nulliparous, involution day 2 and involution day 4 mice when fed vitamin D standard diets (1,000 IU/Kg). Expression of *Cyp2r1* was 1.5-fold lower in involution day 4 mice compared to nulliparous mice (p=0.02; Figure 3B). At the same time, *Cyp27b1* was 1.6-fold lower in involution day 2 mice, compared to nulliparous mice (p=0.007; Figure 3C). In contrast, expression of *Cyp24a1*, involved in vitamin D degradation, was significantly increased during involution day 2 (5.2-fold; p<0.001) and day 4 (2.8-fold; p=0.02), compared to nulliparous mice (Figure 3D).

**Figure 3:**
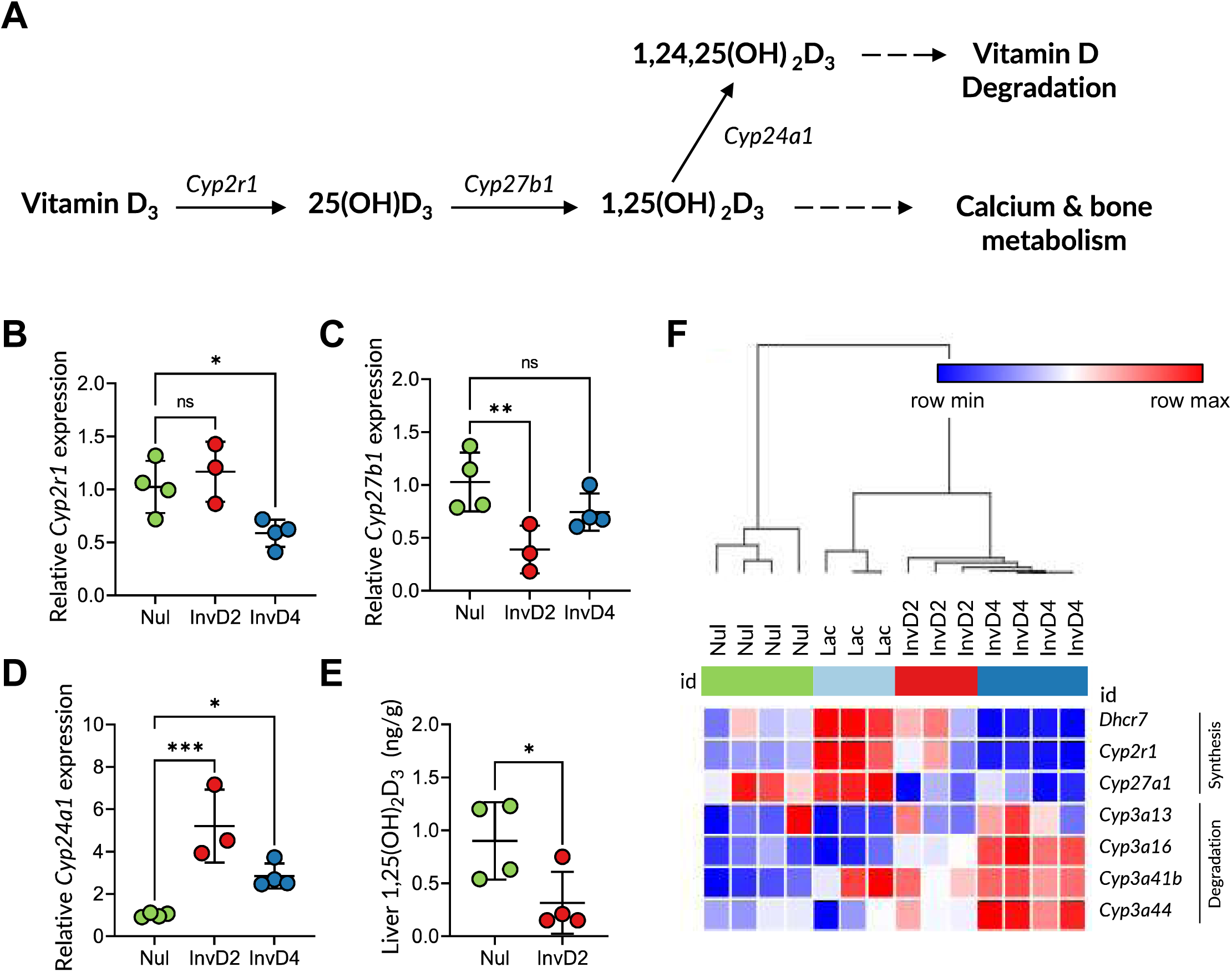
Vitamin D metabolism is impaired in the post-wean liver. (A) Schematic of dietary vitamin D metabolism. (B-F) Livers collected from mice at reproductive time points (nulliparous (nul), involution day 2 (InvD2) and InvD4) were analyzed for vitamin D metabolism pathways using qPCR and RNAseq approaches. Expression of enzymes involved in vitamin D activation (B) *Cyp2r1* and (C) *Cyp27b1*, as well as vitamin D degradation (D) *Cyp24a1* were assessed with qPCR. (E) Tissue-specific levels of active vitamin D metabolite 1,25(OH)_2_D_3_ in the liver of vitamin D supplemented mice, as determined through LC-MS/MS. (F) Hierarchical clustering of genes involved in vitamin D metabolism using an independent murine liver RNAseq data set, reported as mean transcripts per million ±Stdev. P-values are presented, determined through ANOVA with multiple comparisons. ns= not significant; *p<0.05; **p<0.01; ***p<0.001.

To understand the impact of altered gene expression on vitamin D metabolism, nulliparous and involution mice were supplemented with dietary vitamin D (4,000IU/Kg), and the concentration of 1,25(OH)_2_D_3_ in liver tissue assessed. We found a significant reduction in 1,25(OH)_2_D_3_ in involution livers (0.32±0.3ng/g) compared to nulliparous livers (0.90±0.37ng/g; p=0.02; Figure 3E). These findings are consistent with impaired hepatic metabolism during involution. Further, these data suggests that the lack of anti-cancer effects from vitamin D supplementation during this period are due to the lack of the active form of vitamin D.

Using RNAseq data obtained from mouse livers across a reproductive cycle, we next performed a heatmap analysis of the genes involved in vitamin D metabolism. Consistent with our qPCR findings, the heatmap analysis demonstrated expression of genes involved in vitamin D synthesis (*Dhcr7*, *Cyp2r1*, *Cyp27a1*) were reduced during involution. In contrast, genes associated with the degradation of active vitamin D (*Cyp3a13*, *Cyp3a16*, *Cyp3a41b*, *Cyp3a44*) were upregulated (Figure 3F). These observations were replicated in an independent set of mice (Supplementary Figure 1). Together these results suggest suppression of vitamin D activation, combined with active removal of vitamin D in the involution period, leads to reduced serum 25(OH)D and loss of vitamin D anti-cancer activity in our PPBC model.

### 1,25(OH)2D3 treatment post-wean identifies involution as a metabolically vulnerable window

We predicted that treatment of involution mice with active vitamin D (1,25(OH)_2_D_3_) would restore anti-cancer activity. To test this, mice were treated with 1,25(OH)_2_D_3_ (20µg in PBS) on the day of wean, and again on involution day 2. Unexpectedly, all involution treated mice had to be euthanized by involution day 3, due to evidence of acute toxicity (n=7), while vehicle control treated involution mice (n=6) showed no such toxicity (Figure 4A). The toxic effect of 1,25(OH)_2_D_3_ treatment was specific to involution mice, as nulliparous mice treated with 1,25(OH)_2_D_3_ (n=3) or a vehicle control (VC) (n=4) showed no ill-effects from treatment (Figure 4A). Further, nulliparous mice tolerated increased dosing of 1,25(OH)_2_D_3._ We tested treatment in nulliparous mice thrice weekly for 2 weeks, and no evidence of adverse events, including body weight loss or lethargy, were observed.

**Figure 4:**
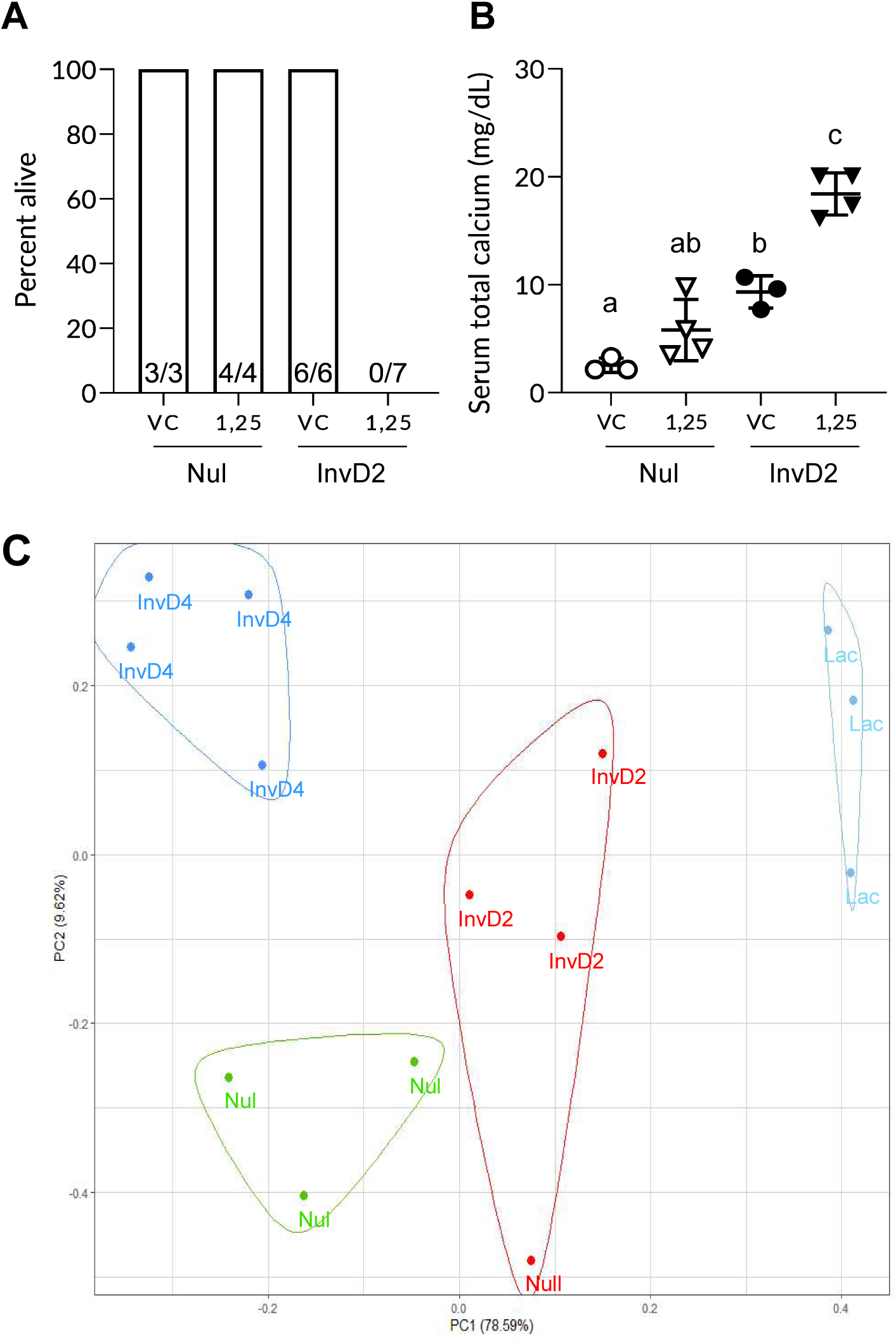
1,25(OH)_2_D_3_ treatment induces toxicity in the involution period, likely due to hypercalcemia. Vitamin D deficiency was established in nulliparous (Nul) and involution day 2 (InvD2) mice through feeding diets deficient in vitamin D content (25IU/kg). On the day of weaning (InvD0), mice were randomized into treatment groups to receive either 1,25(OH)_2_D_3_ (1µg/kg) or vehicle control (5% EtOH in PBS) via subcutaneous injections. In parallel, nulliparous mice were treated. Treatments were re-administered 48 hours later (InvD2), and tissue collected at time of euthanasia. (A) Graph depicting mouse survival following second injection. (B) Serum calcium levels. (C) Principal component analysis (PCA). The expression of 70 murine CYP450 genes were clustered using K-means clustering in R Studio, where k=4. Different colors represent unique clusters. Different letters represent statistical significance determined by ANOVA. Data are mean±Stdev.

Given the role of vitamin D in calcium homeostasis and the toxicity of calcium dysregulation, serum calcium levels were assessed in vehicle control and 1,25(OH)_2_D_3_ treated nulliparous and involution mice. Serum calcium was significantly elevated in involution vehicle control treated mice (9.3±1.5mg/dL), compared to nulliparous vehicle control treated mice (2.5±0.7mg/dL, p=0.01, Figure 4B). This was consistent with our results in mice on chow diets showing baseline calcium is elevated during the window of involution (Figure 2B). Treatment of involution mice with 1,25(OH)_2_D_3_ further elevated serum calcium approximately 2-fold (18.4±2.0mg/dL; p<0.001) to hypercalcemic levels (43). These data demonstrate unique changes in calcium homeostasis following weaning, which is further disrupted by treatment with active vitamin D. No changes in serum calcium were observed in nulliparous mice following 1,25(OH)_2_D_3_ treatment (5.8±2.8mg/dL; p=0.22).

Together, these data identified involution as a vulnerable window with respect to vitamin D metabolism and calcium homeostasis. We next investigated whether the post-wean liver displays gene expression profiles consistent with more extensive alteration in xenobiotic drug metabolism. We performed K-means clustering analysis to evaluate the gene expression patterns of 70 cytochrome P450 (CYP450) genes involved in drug metabolism, selected based on their detectable expression levels in our liver RNASeq dataset (Supplementary Table 2). We found that the expression profiles of CYP450 genes uniquely grouped according to reproductive stage. Specifically, pro-drug activating CYP450 genes were suppressed during involution (Figure 4C). We validated these findings in an independent set of mice (Supplementary Table 1). Together, these data demonstrate a link between reproductive stage and liver CYP450 gene expression. The implication of these results is that the ability of the liver to metabolize xenobiotic drugs is implicate reproductive state dependent.

## Discussion

As breast cancer incidence in young women continues to rise worldwide (44, 45), with young-onset breast cancer carrying a disproportionally high mortality rate (44, 46), the efforts to develop new therapies for young women are becoming increasingly important. Pre-clinical studies have identified weaning-induced mammary gland involution as a pro-tumor inflammatory window of increased breast cancer risk and progression (11–13, 22). In rodents, this postpartum window is amenable to immunotherapeutic intervention (11, 12, 19). Vitamin D has immunomodulatory, anti-cancer properties, and has been suggested as a potential therapeutic agent to reduce risk of young-onset breast cancer (37). Indeed, using a murine model of young-onset breast cancer, we show that vitamin D supplementation reduces tumor growth in nulliparous mice. Further, we find that the anti-tumor mechanism is likely mediated through promotion of an anti-tumor immune milieu. However, in a model of postpartum breast cancer, we find the anti-cancer effects of vitamin D supplementation immediately post-wean are blocked, due to impaired hepatic metabolism of dietary vitamin D to its active form. Attempting to bypass this impaired metabolism, through administering active 1,25(OH)_2_D_3_, proved fatal in post-wean (involution) mice. Toxicity was likely due to dysregulated calcium metabolism post-wean that was exaggerated by treatment with active vitamin D. These results highlight weaning-induced involution as a metabolically vulnerable window of time that needs further exploration in young women. If this post-wean liver biology is confirmed in women, it will be necessary to tailor xenobiotic therapies for young women based on their recent childbirth and lactation history.

One prediction of these murine data is that vitamin D supplementation may be ineffective in women following weaning. While the efficacy of supplementation in this reproductive window remains unknown, it is likely important for maternal health. The majority of research surrounding vitamin D supplementation across the reproductive lifespan focuses on pregnancy and breast-feeding. While previous studies report that vitamin D deficiency in the postpartum window ranges up to ∼72% in US women (35, 36), these studies focus primarily on the lactation window. Further, while there is evidence that vitamin D supplementation during pregnancy and lactation increases serum 25(OH)D (47, 48), the efficacy of vitamin D supplementation post-wean has not been investigated in women. Here, we show that serum levels of vitamin D are deficient in post-wean mice, suggesting increased vulnerability for breast cancer along with other vitamin D related pathologies. Further, vitamin D supplementation during this window is unable to restore serum levels to sufficiency. Critically, as weaning accompanies the return of fertility, our data could suggest that subsequent pregnancies may be at increased risk of vitamin D deficiency despite supplementation. Further understanding of metabolic physiology during involution may be critically important for interpregnancy care. This includes reducing morbidity from vitamin D deficiency associated with common antenatal complications, such as preeclampsia (49).

The suppression in vitamin D metabolism we observed in the post-wean period is likely a direct consequence of weaning-induced liver involution, which we recently reported in rodents (50–52). Relevance to women is suggested, as the human liver also undergoes volume gain with pregnancy followed by post-wean volume loss, consistent with weaning-induced liver involution (52). In mice, liver involution establishes a metastatic niche (50, 51), which may account for increased liver metastasis observed in postpartum breast cancer patients (3, 6, 7). Recently, weaning-induced liver involution has been recognized in clinical guidelines as a risk factor for poor prognosis, and consideration for the care of postpartum patients with a history of solid tumors (53). While the association between liver involution and lactation insufficiency (24), as well as metastatic potential have been described, the effect of liver involution on hepatic metabolism in women remains largely unknown.

In rodents, liver involution is characterized by catabolic metabolism, hepatocyte cell death, stromal remodeling, and pro-tumor immune cell influx (51, 52). Here, we expand on these findings to show that in mice, vitamin D metabolism is suppressed during this post-wean window, rendering vitamin D supplementation ineffective. We show that expression of genes (*Cyp2r1*, *Cyp27b1*) involved in the synthesis of 1,25(OH)_2_D_3_ are reduced post-wean, while a key gene (*Cyp24a1*) involved in the degradation of 1,25(OH)_2_D_3_ is increased. These data suggest overall suppression of vitamin D metabolism to active compound, and the apparent controlled removal of active metabolites. This suggests that tight control of vitamin D metabolites is critical during the involution window. When bypassing this tight regulation of active vitamin D levels by treating with 1,25(OH)_2_D_3_, involution mice became hypercalcemic to toxic levels, necessitating euthanasia. Together these results show that vitamin D concentrations are tightly regulated during the window of involution. This tight regulation is likely due to the unique physiological requirements of weaning induced involution, including increased control over calcium mobilization. Alternatively, the suppression of liver vitamin D metabolism could be an indirect consequence of liver involution, whereby metabolism overall is severely curtailed during the hepatocyte cell death phase of weaning-induced liver involution (51).

Our data suggests that the impact of suppressed liver metabolism post-wean may extend beyond vitamin D activation. We show expression of 70 CYP450 enzymes—enzymes responsible for xenobiotic drug-metabolism—cluster by reproductive stage. These data suggest that the activation and/or elimination of other drugs is likely also to be dependent on reproductive state. Others have reported that pregnancy & lactation induce changes in CYP450 gene expression in women (54, 55). Specifically, activity of CYP2D6 and CYP3A4, key P450 enzymes collectively responsible for the biotransformation of ∼65-75% of all prescribed drugs (56), are increased during pregnancy, and decreased post parturition (54, 55). In contrast, CYP1A, another enzyme that mediates biotransformation of xenobiotic drugs, was reduced during pregnancy, and significantly upregulated after childbirth (55). Our findings extend these pregnancy focused studies by identifying global suppression of xenobiotic metabolism specifically in the post-wean rodent liver.

Our study may have implications for dosing and toxicity of commonly used medicines in the postpartum period. In the context of breast cancer, altered liver metabolism may affect activation of common endocrine (tamoxifen) and chemotherapeutic (cyclophosphamide) agents, which are administered as pro-drugs (57). In a broader context, suppressed P450 activity may alter efficacy of common treatments administered to postpartum women. This includes pro-drugs to treat depression (sertraline)(58) as well as over-the-counter agents, such as antihistamines (loratadine)(57). Further, enhanced degradation of common active drugs, such as those that treat diabetes (metformin)(59) or hypertension (labetalol)(60), might also reduce drug activity. In order to optimize the care and safety of postpartum women, it is essential that future research evaluates the ability of the post-wean liver to metabolize pro-drugs into their active forms, as well as the livers capacity to clear active metabolites. Such studies would ensure a comprehensive understanding of drug metabolism and efficacy during the post-wean period. This information would lead to more accurate dosing and reduce the risk of adverse effects in postpartum women.

Our study adds to the literature regarding vitamin D’s anti-tumor activity in never-pregnant mice. We show that vitamin D supplementation reduced mammary tumor burden through mechanisms that do not rely on inhibition of proliferation, but associate with increased cell death. We found that increased tumor cell death correlated with a reduction in abundance of immature myeloid cells, and increased abundance of mature macrophages. Our data are consistent with other reports that implicate immature myeloid cells as targets for anti-cancer therapy, whereby promoting their differentiation into mature myeloid cells restores immune balance, cytotoxic T cell function, and tumor suppression (12, 39, 61, 62).

Support for the anti-cancer activity of vitamin D being attributable to direct effects on myeloid cell biology comes from prior *in vitro* studies. These studies show that treatment of immature myeloid cells with 1,25(OH)_2_D_3_ prior to co-culture with T-cells reduces their ability to suppress T-cell proliferation (34). Consistent with these studies, we show that vitamin D supplementation associates with relief of T-cell suppression in tumors collected from nulliparous mice, as evidenced by increased expression of T-cell activation/exhaustion markers, GzyB, TOX1, and Ki67. Relief of suppression is likely mediated by vitamin D receptor (VDR) signaling. Previous studies show that VDR knockout in immature myeloid cells increases their T-cell suppression through increased expression of Arg1 and Nos2 (34), markers essential for the suppressive function of immature myeloid cells. However, it is important to note that T-cells also express VDR (63), and treatment with 1,25(OH)_2_D_3_ directly alters T-cell cytokine secretion and skews T-cell polarization (64). Consequently, the potential mechanisms through which vitamin D promotes an anti-tumor immune response may be mediated through either direct or indirect effects on T-cells. Further, as VDR is expressed on most cells of the immune system (63, 65, 66), the role of vitamin D on other immune cells cannot be excluded. Particularly, vitamin D has been shown to have direct effects on B cells (67), dendritic cells (65), neutrophils (66), and natural killer cells (68). In sum, our data broaden the current understanding of vitamin D’s immunomodulatory roles, specifically in breast cancer where the immunomodulatory effects of vitamin D are not as well described.

## Conclusions

We show that dietary vitamin D supplementation is effective at reducing growth of mammary tumors that arise in young, nulliparous mice. This anti-tumor effect is achieved through mechanisms independent of tumor cell proliferation. Instead, the anti-tumor effect of vitamin D supplementation associates with macrophage maturation, T-cell activation, and caspase-dependent tumor cell death. These results confirm previous studies implicating vitamin D as an anti-tumor immunotherapeutic agent. Unfortunately, dietary vitamin D supplementation proved ineffective during involution due to lack of metabolism to active form. Our findings emphasize the metabolic vulnerability of the postpartum period of involution. The suppression of liver xenobiotic metabolism we identified post-wean has implications for drug efficacy and safety in postpartum women. Specifically, the dietary form of vitamin D (i.e., a pro-drug) was ineffective, while 1,25(OH)_2_D_3_ (i.e., the active form) supplementation led to hypercalcemia and mortality. Further studies are warranted to assess the ability of the human liver to metabolize commonly prescribed xenobiotic drugs in the post-wean period. If confirmed in women, this sex-specific liver biology has broad implications for maternal health, as young women may be highly resistant to pro-drug efficacy and could be more vulnerable to toxicity.

## Methods

### Cell culture

The mouse mammary cancer cell line, D2A1, was donated by Ann Chambers (University of Western Ontario, London, Ontario, Canada), and cultured in High Glucose DMEM with L-glutamine, supplemented with 10% FBS (S11550H, R&D Systems). For the tumor injection study, D2A1 cells were resuspended at 1×10^6^ cells/mL in 1×PBS. Final cell injects used 20,000 D2A1 cells in 20μl PBS. Cells were confirmed to be of proper cell origin and mycoplasma free (July 2022; Idexx Bioresearch, Columbia, MO, USA).

### Animal studies

#### Ethics Approval

Ethics approval was obtained from the Oregon Health & Science University (Approval number TR01_IP00000967). All mice were obtained from Charles River Laboratories (Wilmington, MA, USA).

#### Breeding

To obtain specific reproductive stages (including involution days 2, 4, and 6), female BALB/c mice of 10-12 weeks old were bred with male C57Bl/6 mice of 12 weeks old. This breeding scheme induces fetal-maternal tolerance mechanisms, modelling those present in humans (13). Two days post-parturition, pup number was normalized to 6-8 pups/dam to ensure equal lactational load. Following 9-13 days of lactation, pups were removed to initiate synchronized involution.

#### Diets to model Vitamin D Deficiency or Supplementation

At the time of breeding, mice were started on custom diets, which contained deficient (25IU/kg; TD.210353, Envigo) or supplemented (4,000IU/kg; TD-210354, Envigo) levels of vitamin D. Mice were maintained on these diets for the duration of the study.

#### Tumor Implantation

Mice were anaesthetized using isoflurane, and murine mammary cancer cells (D2A1, 2×10^4^ cells in 20µL PBS) were inoculated into the fourth inguinal left and right fat-pad of involution day 2, and of age-matched nulliparous mice. Tumor growth was tracked twice a week, for 4 weeks.

#### Active Vitamin D treatment

At the time of weaning (involution day 0), mice were randomized into treatment groups, and treated with either (i) 1α,25(OH)_2_D_3_ (1µg/kg, H0891ML, Sigma-Aldrich) or (ii) vehicle control (5% EtOH in PBS), with drug delivered by subcutaneous injection. In parallel, nulliparous mice were also treated. Treatments were repeated 48 hours later (involution day 2), and tissue collected at time of euthanasia (approximately 8 hours after the final injection).

#### Tissue Collection and Euthanasia

Mice were euthanized at specific reproductive stages (nulliparous, and involution days 2, 4, and 6) or at tumor endpoint. Total blood was collected by cardiac puncture under anesthesia (Isoflurane, USP; cat no. 66794-017-25; Piramal Critical Care), and mammary tumor, liver and hind limbs were collected. For breast cancer studies, tumors were dissected and processed for immunohistochemistry analysis. For liver analysis, tissue was snap-frozen in liquid nitrogen for LC-MS/MS analysis.

### Gene expression analyses

#### Quantitative PCR

Livers were collected from mice at the following reproductive stages: nulliparous (n=4), involution day 2 (n=3) and involution day 4 (n=4). RNA was isolated from livers, as described in (52). Total cDNA was reverse transcribed from 500ng of RNA using iScript IV VILO cDNA synthesis kit (Bio-Rad), with reactions incubated at 25°C for 5 min, 42°C for 25 min, and 85°C for 5 min. Quantitative PCR (qPCR) was performed using a CFX Opus 384 (Bio-Rad) Real-Time PCR systems. Primer pairs were obtained from Integrated DNA Technologies and are detailed in Supplementary Table 1. Reactions were performed in a total volume of 10ul, containing 1×SYBR green master mix (Bio-Rad), 4µM forward and reverse primers, and cDNA template.

#### K-means clustering

Clustering analyses were performed using normalized RNAseq data (GEO accession GSE260714). K-means clustering focused on gene expression profiles from livers dissected from mice at reproductive stage: nulliparous (n=4), lactation day 10 (n=3) and involution day 2 (n=3) & involution day 4 (n=4). K-means clustering was used to categorize expression of 70 cytochrome P450 genes (Supplementary Table 2) into distinct clusters. The number of clusters (k) was defined as 4, based on the number of reproductive groups analyzed (Nulliparous, lactation, involution days 2 & 4). Data are presented using PC1 and PC2.

#### Heatmap analyses

Heatmap analyses were performed using normalized RNAseq data. Gene expression in livers collected from mice at reproductive stages (nulliparous, lactation day 10, involution days 2 & 4) was clustered based on the expression of vitamin D metabolic genes (degradation: *Cyp3a13*, *Cyp3a16*, *Cyp3a41b*, *Cyp3a44*); synthesis (*Dhcr7*, *Cyp2r1*, *Cyp27a1*)). Hierarchical clustering was performed using Morpheus (69), an online tool for data analysis and visualization, grouping columns by reproductive stage.

### Serum analysis

Total serum calcium was measured using a colorimetric calcium assay kit (Cell Biolabs, MET-5121) as per the manufacturer’s instructions.

### LC-MS/MS analysis

#### Sample Preparation and Calibrator Samples

The Chemicals and Reagents used are described in the Supplementary Methods. Liver tissue samples were homogenized at 1 gram/3 ml PBS using Omni International Bead Rupter with 2 cycles 30 sec 5 m/s with a 5 second pause between cycles. Calibrators were prepared using commercial stock solutions that were diluted to working stock solutions as required and stored at −80°C in methanol. The calibration curve used for measurement in tissue or plasma samples was prepared in serum substitute matrix, consisting of 5% bovine serum albumen in 0.9% saline. Calibrants were spiked with <5% of working stock solutions and shaken for 30 minutes at room temperature in the dark. Unknown samples or calibrants (100 μl) were transferred to fresh Eppendorf tubes and 0.05 ng each internal standard was added in 10 μl methanol. Samples were shaken for a further 15 minutes in the dark. Disassociation solvent was added (300 μl of 1:1 water:isopropanol v:v with 1% formic acid) and samples were vortexed vigorously for 5 minutes on a pulsing vortexer at 2500 rpm, before an additional 15 minutes of shaking in the dark to complete the disassociation. Samples were then centrifuged to pellet any debris and supernatant was loaded onto 400 μl SLE+ cartridges and allowed to absorb for 5 minutes (no positive pressure was used to load). After 5 minutes, 1 ml of heptane is added to SLE+ cartridge and allowed to elute by gravity (this step was performed twice). Elution was completed by a short burst of positive pressure.

Prior to loading of samples onto SLE+ cartridge, PTAD solution (200 μl) is added to the elution tubes. PTAD solution is freshly prepared for each use in two stages. First PTAD is dissolved at 5 mg/ml in ethyl acetate in a foil covered tube to protect from light, then diluted with heptane to a final concentration of 1 mg/ml and stored on ice in a foil covered tube until use. Tubes with eluted sample and PTAD solution are capped, briefly vortexed to mix sample and shaken for 1 hour in the dark at ambient. Samples are dried for 30 minutes at 40°C using 1L/min in a Biotage TurboVap unit and reconstituted in 100 μl buffer (70:30:0.05 methanol:water:methylamine v:v:v), vortexed and transferred to vials for LC-MS/MS sample analysis.

#### LC-MS/MS Analysis of Vitamin D and metabolites

Vitamin D and metabolites were analyzed using a 5500 Q-TRAP hybrid/triple quadrupole linear ion trap mass spectrometer (AB SCIEX, Framingham, MA) with electrospray ionization (ESI) operating in the positive mode. The mass spectrometer was interfaced to a Shimadzu (Columbia, MD) SIL-20AC XR auto-sampler followed by two LC-20AD XR LC pumps. The instrument was operated with the following settings: source voltage 5500 kV, GS1 40, GS2 40, CUR 40, TEM 500 and CAD gas MEDIUM. The scheduled Multiple Reaction Monitoring (MRM) transitions presented in the table below were monitored with an 80 second window. Each compound was infused individually and instrument parameters optimized for each MRM transition. The high performance liquid chromatography (HPLC) mobile phase consisted of two solvents, Buffer A (water with 0.1% formic acid and 60 μl/L of methylamine), Buffer B (methanol with 0.1% formic acid and 60 μl/L of methylamine). The HPLC column used was a Phenomenex Kinetex EVO C18 50x.2.1 (Torrance, CA). The HPLC gradient was at 0.5 ml/min flow rate as follows: 50% B for 0.5 minutes, gradient to 65% B over 2 minutes, 65% B for 3 minutes, gradient to 98% B over 2 minutes, 98% B for 2.5 minutes, gradient to 50% B over 0.5 minutes, 50% B for 1.5 minutes, with total run time of 13 minutes. MRM Transitions used, lower limit of detection (LLOD), lower limit and upper limit of Quantitation (LLOQ/ULOQ) are provided in Supplementary Table 3.

### Bone mineral density analysis

Left hind limbs were collected from mice and stored in 70% EtOH. Dual-energy x-ray absorptiometry (DXA) scans and analysis were performed *ex vivo* using a Faxitron UltraFocus DXA (Hologic). Bone mineral density (BMD) of each sample was quantified in the proximal tibia, in the region of the bone extending 3mm below the growth plate.

### Immunohistochemistry & multiplex immunohistochemistry

Immunohistochemistry (IHC) and multiplex IHC were performed on 4μm thick sections of formalin-fixed, paraffin-embedded tissue samples. Slides were de-waxed by incubating at 60°C for 1 hours, prior to passing through xylene and gradient ethanol (100%, 95%, 70%, 50%) then water. Antigen retrieval was performed in a pressure cooker at 115°C for 20 min, using either low (Citrate, Dako, pH 6) for Ki67 and multiplex panels; or high pH (EDTA, Biocare, pH 9.0) for cleaved caspase 3 (CC3) staining.

For immunohistochemical analysis, slides were incubated with primary antibodies, Ki67 (Cell Signaling, 1:400, overnight at 4°C), or Cleaved Caspase-3 (CC3; Cell Signaling, 1:150, 1 h at room temperature). Slides were washed thrice in TBST prior to incubation with a pre-diluted HRP-conjugated secondary antibody (Nacalai USA Histofine). Antibody binding was visualized using DAB peroxidase (HRP; Vector Laboratories), as per manufacturer’s instructions. Slides were counterstained with hematoxylin prior to dehydrating and coverslipping. Stained tissue sections were imaged using an Aperio Scanscope (Leica Biosystems), and staining was quantified using Aperio ImageScope software (Leica Biosystems).

For multiplex analysis, after antigen retrieval, slides were cyclically probed as described previously (12, 70) in the following order with the indicated antibody, dilution and incubation times: Myeloid Panel: Cycle 0 Hematoxylin (Agilent, Ready-to-use), Cycle 1 GR-1 (Invitrogen, 14-5931-82, 1:100, 1hr), Cycle 2 F4/80 (Cell Signaling, 70076S, 1:500, 1hr), Cycle 3 CD45 (BD Pharmingen, 550539 (Clone 30F11), 1:50, 1hr), Cycle 4 CD11b (Abcam, ab133357, 1:30,000, 1hr). Lymphoid Panel: Cycle 0 (Hematoxylin (Agilent, Ready-to-use), Cycle 1 CD4 (Cell Signaling, 25229, 1:50, overnight), Cycle 2 CD45 (BD Pharmingen, 550539 (Clone 30F11), 1:50, 1hr), Cycle 3 Ki67 (Cell Signaling, 12202, 1:800, 1hr), Cycle 4 Tox1 (Abcam, ab237009, 1:300, 1hr), Cycle 5 CD3 (Abcam, ab16669, 1:100, 1hr), Cycle 6 (Granzyme B, an4059, 1:500, 1hr), Cycle 7 (eBiosciences, 14-0195-85, 1:100, 1hr). Detection of primary antibodies were performed with either anti-rabbit or anti-rat horseradish peroxidase polymers (Histofine Simple Stain MAX PO, Nacalai USA) followed by detection with peroxidase substrate 3-amino-9-ethylcarbazole (AEC). Stains were scanned using Aperio ImageScope AT2 (Leica Biosystems) at 20× magnification. Image processing and analysis were performed as previously reported (12, 70) utilizing CellProfiler (Broad Institute) and FCS-Express software (DeNovo Software, Glendale, CA).

### Flow Cytometry

Flow cytometry data was obtained as previously described (24). Briefly, mammary glands were collected from mice on chow diets (∼1000IU/kg vitamin D content) at reproductive stages: nulliparous, involution day 2 and involution day 4. Digested mammary glands were blocked with CD16/32 (eBioscience, 1:100) and stained with Live Dead (aqua, Invitrogen, 1:500) for 30 min. Cells were stained with for CD45 (30-F11, 1:800, Pe-Cy7); CD11b (M1/70, 1:200, BV711); F480 (BM8, 1:400, APC); Ly6C (HK1.4, 1:400, PerCP-Cy5.5); Ly6G (1A8, 1:400, APC-Cy7) for 30 min at room temperature. Cells were fixed (BD Cytofix) and run on an 18-color flow cytometer (Fortessa BD). Data were analyzed using FlowJo software.

### Statistical analyses

Statistical analyses were performed using Graph Pad Prism (Version 9.2.0). Differences between two treatment groups were assessed using Two-Tailed T-Tests; while differences between >2 treatment groups were assessed using analysis of variance (ANOVA) with post-hoc comparisons performed. Data were considered significant when p<0.05. All data is presented as mean±Standard Deviation (StDev).

## Acknowledgements

We would like to thank AeSoon Bensen for her assistance with animal experimentation and data collection. We would also like to thank Weston Anderson for their assistance editing the manuscript.

## Author contributions

Experimental conception and design were conducted by Bernhardt SM, Schedin P. Data collection was performed by Bernhardt SM, Ozaki M, Zhang Y, Bleyle L, De Wilde E, Fornetti J, Betts C. Data analysis was performed by Bernhardt SM, Zhang Y, De Wilde E, Fornetti J, DeBarber A, Liberty A, Xia Z, Schedin P. The draft of the manuscript was written by Bernhardt SM, Liberty A, Schedin P, with all authors providing intellectual input into the final manuscript. All authors have read and agreed to the published version of the manuscript.

