## Supplementary Files for "Altered liver metabolism post-wean abolishes efficacy of vitamin D for breast cancer prevention in a mouse model"

### Supplementary Methods

Acetonitrile, methanol, isopropanol and water were LC-MS grade from Honeywell - Burdick and Jackson (Muskegon, MI). ACS grade formic acid, Bovine serum albumen (BSA), Phosphate Buffered Saline (PBS) 1X pH 7.4 were obtained from Thermo Fisher Scientific (Fair Lawn, NJ). Derivatization reagent 4-phenyl-1,2,4-triazoline-3,5-dione (PTAD), methylamine solution, ethyl acetate and sodium chloride were from Sigma-Aldrich (Chicago, IL). Vitamin D2, Vitamin D3 and metabolite reference standards were from Cerilliant (Round Rock, TX). Deuterated internal standards were from Cambridge Isotope Laboratories (Tewksbury, MA). Biotage 400 l SLE+ (Supported Liquid Extraction+) cartridges were from Biotage (Uppsala, Sweden). Soft tissue homogenizing mix 1.4 mm ceramic beads (used with 2 ml soft tissue tubes) from Omni International (Kennesaw GA).

### Supplementary Tables

**Supplementary Table S1: Primer sequences for quantitative PCR against mouse genes**

| Gene | Forward | Reverse |
|---|---|---|
| *Cyp24a1* | 5'-CTGGCCTGGGACACCATTT-3' | 5'-CTCCGTGACAGCAGCGTACA-3' |
| *Cyp27b1* | 5'-CAGAGCGCTGTAGTTTCTCATCA-3' | 5'-CGTTAGCAATCCGCAAGCA-3' |
| *Cyp2r1* | 5'-AAACTACAACCAATGTGCTCCG-3' | 5'-ATTCCCAAGAAGGTCTCCTGT-3' |
| *Gapdh* | 5'-TCAACAGCAACTCCCACTCTTCCA-3' | 5'-ACCCTGTTGCTGTAGCCGTATTCA-3' |

**Supplementary Table S2: List of murine CYP450 genes analyzed in principal component analyses.**

| | | | |
|---|---|---|---|
| Cyp17a1 | Cyp2c39 | Cyp2j5 | Cyp4a14 |
| Cyp1a1 | Cyp2c40 | Cyp2j6 | Cyp4a31 |
| Cyp1a2 | Cyp2c44 | Cyp2j9 | Cyp4a32 |
| Cyp1b1 | Cyp2c50 | Cyp2r1 | Cyp4b1 |
| Cyp20a1 | Cyp2c54 | Cyp2s1 | Cyp4f13 |
| Cyp26a1 | Cyp2c55 | Cyp2u1 | Cyp4f14 |
| Cyp26b1 | Cyp2c67 | Cyp39a1 | Cyp4f15 |
| Cyp27a1 | Cyp2c68 | Cyp3a11 | Cyp4f16 |
| Cyp2a12 | Cyp2c69 | Cyp3a13 | Cyp4f17 |
| Cyp2a22 | Cyp2c70 | Cyp3a16 | Cyp4f18 |
| Cyp2a4 | Cyp2d10 | Cyp3a25 | Cyp4f39 |
| Cyp2a5 | Cyp2d22 | Cyp3a41a | Cyp4v3 |
| Cyp2b10 | Cyp2d26 | Cyp3a41b | Cyp51 |
| Cyp2b13 | Cyp2d40 | Cyp3a44 | Cyp7a1 |
| Cyp2b9 | Cyp2d9 | Cyp3a57 | Cyp7b1 |
| Cyp2c29 | Cyp2e1 | Cyp3a59 | Cyp8b1 |
| Cyp2c37 | Cyp2f2 | Cyp46a1 | |
| Cyp2c38 | Cyp2g1 | Cyp4a10 | |

**Supplementary Table S3: Precursor ion masses are derivatized-methylamine complexes, RT is retention time and IS is internal standard.**

| Compound ID | RT (min) | Precursor m/z | Fragment m/z | DP | CE | CXP | EP | LLOD | LLOQ | ULOQ | IS Used |
|---|---|---|---|---|---|---|---|---|---|---|---|
| **1,25 di(OH) D3** | 4.8 | 623.4 | 314.1 | 40 | 30 | 15 | 8 | 0.01 | 0.05 | 50 | d3-1,25 di(OH)D3 |
| **1,25 di(OH) D2** | 5.2 | 635.4 | 314.1 | 40 | 30 | 15 | 8 | 0.01 | 0.05 | 50 | d3-1,25 di(OH)D3 |
| **25(OH) D3** | 6.8 | 607.4 | 298.1 | 50 | 30 | 10 | 8 | 0.005 | 0.01 | 50 | d6-25(OH) D3 |
| **25(OH) D2** | 7.0 | 619.4 | 298.1 | 50 | 25 | 10 | 8 | 0.005 | 0.01 | 50 | d3-25(OH) D2 |
| **Vitamin D3** | 8.5 | 591.4 | 298.1 | 50 | 25 | 10 | 8 | 0.005 | 0.01 | 50 | d3-Vitamin D3 |
| **Vitamin D2** | 8.5 | 603.4 | 298.1 | 50 | 25 | 10 | 8 | 0.005 | 0.01 | 50 | d3-Vitamin D3 |
| **24,25di(OH) D3** | 4.0 | 623.4 | 298.1 | 50 | 35 | 10 | 8 | 0.005 | 0.05 | 50 | d3-1,25 di(OH)D3 |
| **d3-1,25 di(OH) D3(IS)** | 4.8 | 626.4 | 317.1 | 40 | 30 | 15 | 8 | n/a | n/a | n/a | n/a |
| **d6-25(OH) D3(IS)** | 6.8 | 613.4 | 298.1 | 50 | 30 | 10 | 8 | n/a | n/a | n/a | n/a |
| **d3-25 (OH) D2(IS)** | 7.0 | 622.4 | 301.1 | 50 | 25 | 10 | 8 | n/a | n/a | n/a | n/a |
| **d3-Vitamin D3(IS)** | 8.5 | 594.4 | 301.1 | 50 | 25 | 10 | 8 | n/a | n/a | n/a | n/a |

### Supplementary Figures

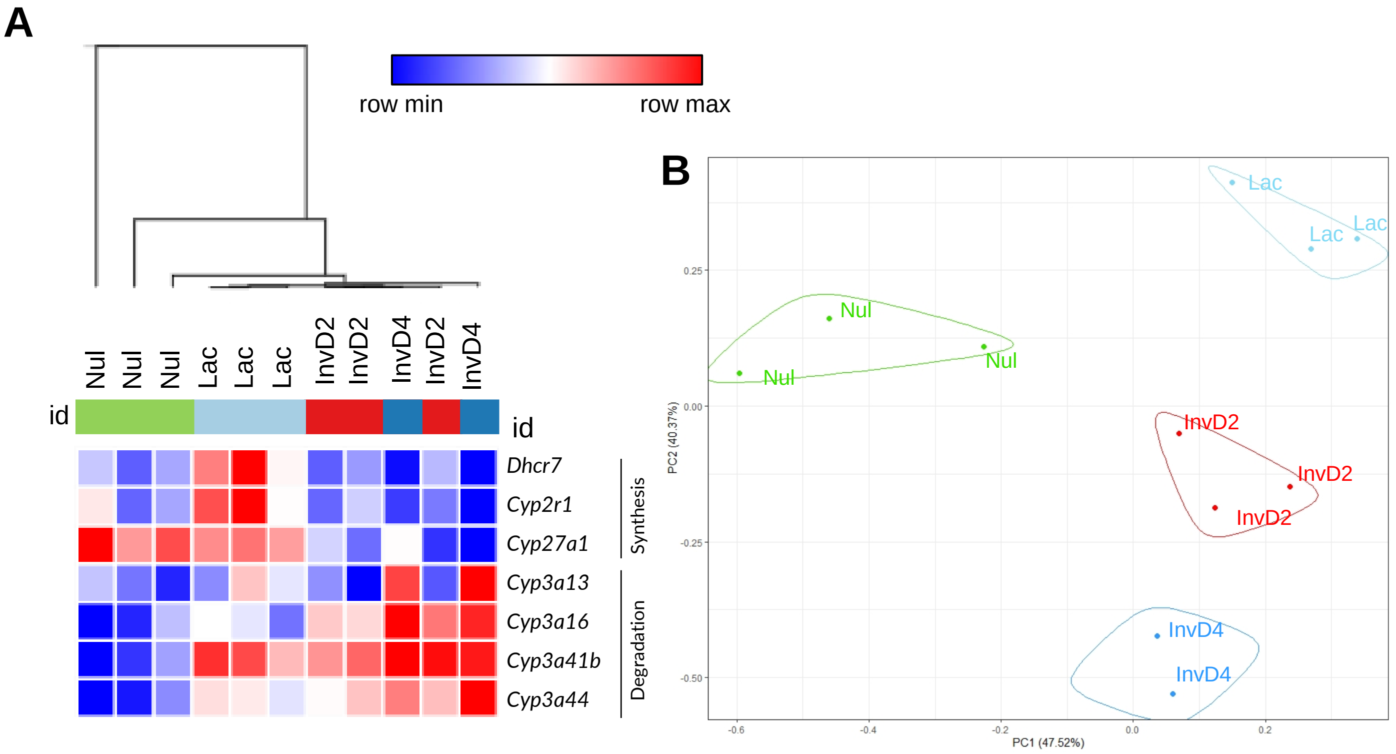


**Supplementary Figure S1: Validation of altered liver metabolism across a reproductive cycle.** RNAseq experiments were repeated using a second batch of mice (purchased from Jackson Laboratory). Livers were collected from adult Balb/C mice (n=2-3 mice/group) on vitamin D standard diets (1,000IU/kg), at reproductive time points: Nulliparous (never-pregnant; Nul), lactation day 10 (Lac), involution day 2 (InvD2) & day 4 (InvD4). RNAseq was performed on whole liver tissue. (A) Hierarchical clustering of genes involved in vitamin D metabolism. (B) Principal component analysis (PCA). The expression of 70 murine CYP450 genes were clustered using K-means clustering in R Studio, where k=4.
